# Re-evaluation of homozygous lethality for EF1A indel polymorphism in *Pisaster ochraceus*

**DOI:** 10.1101/584235

**Authors:** John P. Wares, Paige J. Duffin

## Abstract

A genotypic polymorphism in the sea star *Pisaster ochraceus* has been associated with possible overdominant maintenance of diversity, and subsequent studies of this polymorphism suggested that intermittent disease outbreaks could be a driving factor in this system. However, comparative transcriptomic studies of individuals carrying distinct genotypes at the elongation factor 1-alpha (EF1A) region indicated that the marker was not accurately describing the constitutive differences among individuals. Here we more thoroughly assess this EF1A intron region to better understand how polymorphic diversity could be associated with differential disease outcomes and physiological responses, and find that the underlying genetic model is incorrect. In fact, rather than an instance of homozygous lethality, it is clear that previous genotyping efforts were misled by a PCR artefact. We reanalyze results from two previous studies to show that the effects are not as clear as believed.

## Introduction

The association of genomic diversity with biologically relevant phenotypes remains an elusive problem for most organisms. To identify the contribution of a particular genotype to a trait typically requires some control over parentage and kinship of the studied individuals; for example, the economic importance of trout hatcheries enabled relatively early identification of quantitative trait loci associated with spawning time (Sakamoto et al. 1999) and thermal tolerance (Jackson et al. 1998) in the rainbow trout, *Oncorhynchus mykiss*. When traits can only be studied from individuals in natural populations without this kind of control over mating, association studies may allow discovery of candidate loci to predict traits or the potential for adaptation (Bonin et al. 2006, Hamlin & Arnold 2015). As this latter type of variation and its relationship with phenotype is often inferred rather than directly tested, it is key to understand and evaluate the underlying assumptions that are critical to this inference.

Consideration of the specificity of genomic diversity for behavioral or physiological traits are often more difficult to ascertain. Examples include particular genotypes that appear to be disease tolerant (Vollmer & Kline 2008), or allelic or genotype diversity that predicts parasite load (Brock et al. 2015, Montano-Frías et al. 2016). Classically, our understanding of the fitness consequences of genomic diversity is associated primarily with the avoidance of inbreeding depression, i.e. more diverse populations are less susceptible to developmental and disease problems (Hedrick 1995, Brock et al. 2015). Strong environmental challenges also leave signatures in genome-wide scans (Campbell-Staton et al. 2017, Schiebelhut et al. 2018), with small numbers of markers indicating significant and likely causal shifts in frequency. It is in these instances that researchers have started to identify ecologically-important genes (EIG; Skovmand et al. 2018), where the genotype has likely influence on function or phenotype.

However there are multifactorial reasons for these relationships: is it pertinent to the location of the individual being sampled, or the environmental background in which it has survived and developed (Ewers-Saucedo et al. 2016)? Statistical inference of these genotype-phenotype patterns will often have a tremendous amount of noise because any single marker will be found in highly diverse genomic backgrounds (Stuart et al. 2017, Weber et al. 2017). Attention to the particular individuals that suggest additional complexity to the model is warranted to ensure that the baseline assumptions of the inference are being met.

In recent years, a devastating pandemic has decimated populations of sea stars (Echinodermata: Asteroidea) in diverse coastal habitats, most notably along the west coast of North America (Hewson et al. 2014, Eisenlord et al. 2016, Menge et al. 2016). The search for information that could guide our understanding and management of this outbreak, called “sea star wasting disease” (SSWD), was rapid and involved diverse research teams. Some focused on likely causes (Hewson et al. 2014), with results that were more certain for some taxa than others. Some excellent instances involved pre-exposure contrasts with post-SSWD populations, highlighting genomic regions that appear to be determinant of survival (Schiebelhut et al. 2018). Others focused on the potential for EIGs to indicate that diverse populations would harbor heritable diversity in resistance or tolerance to this syndrome. Wares & Schiebelhut (2016) showed that across diverse locations, the incidence of SSWD was lower in individual *Pisaster ochraceus* that harbored one of two presumed genotypes at the elongation factor 1-alpha (EF1A) locus. Briefly, the genotype had been determined by electrophoresis on a high-percentage agarose gel used to distinguish alleles separable by a 6-bp gap in an intron of EF1A; Pankey & Wares (2009) generated results that appeared to show that one genotype was consistently missing and likely lethal when homozygous.

The apparent relationship between these genotypes and stress tolerance was supported by a temperature challenge study that evaluated RNA transcript abundance in *P. ochraceus* of both recognized EF1A genotypes under ambient and temperature-stressed conditions (Chandler & Wares 2017). This study had three important outcomes: the first was that individuals with distinct recognized genotypes at the EF1A marker had significantly distinct patterns of transcript expression under ambient conditions, and the second was that when heat-stressed, individuals of the ‘tolerant’ genotype had far less transcriptional response than individuals of the other gel-based genotype. However, the third was that the results of this study were partially sensitive to the inclusion or categorization of one of the 10 individuals in the study. Individual ‘*Po*5’ (a ‘homozygote’) exhibited RNA transcript levels at ambient and elevated temperatures that suggested a transcriptional phenotype more similar to the heterozygous individuals in the study, prompting discussion that the causal mutation could be tightly linked to the indel polymorphism that was being screened.

Here we explore this heterogeneity among individuals in the Chandler & Wares (2017) study in greater detail, focusing on what linked genomic diversity in the ‘outlier’ individual *Po5* could be of greatest import for further analyses of this species in the context of disease survival. As the diversity represented by the EF1A genotype seems to have true transcriptional phenotype consequences (Chandler & Wares 2017) and potentially ecologically important effects (Wares & Scheibelhut 2016), a better understanding overall of the diversity in this gene region will guide exploration of physiological phenotypes among individuals that have survived SSWD. What we show here, using diverse approaches with DNA sequencing, more direct methods for genotypic analysis, and physiological assays, is that the underlying model of inheritance at the EF1A marker inferred by Pankey & Wares (2009) and applied to subsequent analyses (Wares & Schiebelhut 2016, Chandler & Wares 2017) is incorrect, rendering the prior conclusions uncertain. Our reanalysis of data from previous studies clarifies our understanding of the ecological relevance of this gene region.

## Methods

### Sequencing of EF1A intron

All individual *Pisaster ochraceus* from the RNA differential expression study of Chandler & Wares (2017) were sequenced for this re-analysis – since it was the diversity in this fragment used to categorize the RNA expression data in that paper – along with congeners *P. giganteus* and *P. brevispinus*. The individual *P. ochraceus* were collected in 2016 from the Friday Harbor Laboratories marine reserve (Chandler & Wares 2017); the congeneric tissues were made available by I. Hewson.

New EF1A forward primer (5’-GACAACGTTGGTTTCAACGTGAAGAACG-3’) was developed from previous sequence data from the upstream coding region (Pankey & Wares 2009); the reverse primer for amplification of the complete intron region is EF2-3’ from Palumbi (1996). We amplified the entire EF1A intron region from which Pankey & Wares (2009) developed their marker, using strategies to minimize PCR-mediated recombination (Lahr & Katz 2009), including minimal template concentration and use of Phusion High-Fidelity polymerase (Thermo Fisher) with a 90 second extension time and a 65°C annealing temperature. PCR amplicons were cleaned with a standard exonuclease protocol (Wares et al. 2009) prior to cloning and sequencing.

Blunt end ligation and cloning was performed using an Invitrogen Zero Blunt TOPO PCR cloning kit for sequencing with the rapid transformation protocol; transformants were then PCR amplified with above primers, treated with exonuclease as above, and Sanger sequenced from both ends to obtain greatest direct haplotype lengths with high accuracy, since the amplified region is ~1kb.

Sequences were edited in CodonCode Aligner v6.0.2, with site quality scores <20 considered ambiguities. For individuals with multiple cloned sequences, reads were aligned to determine distinct alleles separable by one or more observed mutations, for which there had to be confirmation in more than one sequenced read. A consensus sequence was generated for each allele in each individual. All consensus sequences were aligned using the local alignment settings in Aligner, and re-evaluated using the local alignment tools in Geneious R11.1.5; a neighbor-joining phylogram (HKY) was generated in Geneious using the congeneric sequences to root the tree. Evaluation of polymorphisms in the intron region were characterized for sequences with and without the insertion-deletion mutation used to classify individuals in prior studies; analysis of nucleotide diversity and Tajima’s D were calculated using the popGenome package in R (Pfeiffer et al. 2014).

### Capillary electrophoresis of EF1A marker

To clarify the individual genotypes specific to the 6-bp insertion/deletion mutation in the intron of EF1A in *P. ochraceus*, the shorter region characterized in Pankey & Wares (2009) and used in Wares & Schiebelhut (2016) was re-amplified for all individuals from Wares & Schiebelhut (2016) as well as new specimens collected in 2018 (see below). Rather than genotyping on 2% agarose, we used a Fragment Analyzer capillary electrophoresis system (Georgia Genomics Facility). The interpretation of electrophoretograms followed a set of objective rules which were used to characterize the allelic peaks. Because the two focal alleles differ by only 6-bp, software-automated allele calling was insufficient (for example, in heterozygotes, one of the two peaks was often not recognized/labeled), all individuals were reviewed manually. There was enough inherent variation among peak sizes such that alleles were characterized by a small range of expected bp sizes. The variation among product peak sizes is likely due, in part, to the manner in which samples were handled using the Fragment Analyzer system, because we noticed that variation within samples of a given row (of a 96-well plate) was anecdotally lower than that of the entire sample set. Therefore, we made sure each row contained at least one heterozygous individual to help calibrate the analyses with the expected bp sizes of each allele in order to more accurately assign homozygous genotypes to electrophoretograms displaying a single peak.

### Genotypic analysis

In Wares & Schiebelhut (2016), we followed the inference from Pankey & Wares (2009) that for the insertion-deletion mutation there were only two genotypes: a homozygote for the smaller fragment, or an individual heterozygous for the two fragments. Following more extensive cloning and capillary electrophoresis (above), it became clear that all three genotypes for that polymorphism are extant. We therefore re-analyze the survivorship data from Wares & Schiebelhut (2016) to identify whether any one genotype is distinctive from the others regarding their frequency of survival. In that paper, the biological effect was considered to be the difference in proportion of “homozygous” individuals at this marker that appeared symptomatic, as compared to the same proportion appearing among heterozygous individuals, such that a positive value suggested heterozygotes were either more tolerant of or resistant to SSWD. Here, we simply assess at each location the proportion of each genotype that were symptomatic to identify if there are clear differences. As before (Wares & Schiebelhut 2016) we test this hypothesis with a GLM incorporating random effects, location, and genotype on disease status.

### Collection of individuals (2018)

Individual *P. ochraceus* (n=18) were again collected in 2018 from the Friday Harbor Laboratories marine reserve with written permission from the Scientific Director (Professor Megan Dethier). All *P. ochraceus* in this study were collected on June 14, 2018 from the intertidal, each at approximately MLLW, from just north of Point Caution, San Juan Island, Washington (48.557°N, 123.017°W). Each individual was weighed (wet mass, g) and ~10μg of tube feet were excised with sterile scalpel blade for initial genotyping as in Wares & Schiebelhut (2016).

Tissues for genotypic work were placed into 100μl of 10% w/v Chelex-100 and nucleic acids stabilized using the protocol of (Casquet et al. 2012). These samples were then assayed using the PCR conditions of Wares & Schiebelhut (2016) to determine electrophoretic phenotype; they were later re-genotyped using the methods noted above for the same PCR fragment. All but one collected individual were returned in good condition to the intertidal following experimental work.

### Laboratory acclimation and experiment

Individual stars were maintained in flow-through sea tables (~0.5L/min flow) with ambient fluctuations in water temperature for 3 days with *ad libitum* bivalve diet; they were then placed into individually-numbered 5mm mesh bags (“pecan bags”) for the duration of the experiment. They remained fasting in the ambient-fluctuation environment for 48 hours (as in Fly et al. 2012) before movement to the Ocean Acidification (OA) laboratory at FHL for the stress treatment.

In the OA lab, we were able to provide unfiltered seawater at constant set temperature for the duration of the experiment. Water temperature was evaluated regularly for pH, nitrates, and temperature using a YSI meter and Onset Hobo Tidbits in representative chambers. Each *P. ochraceus* was maintained in an individual 3L flow-through seawater chamber (0.1 L/min); chambers were maintained at steady temperature within larger flow-through coolers and individually drained, so no waste or exudate from any single star could influence any other individual. Water changes were performed daily for each individual chamber. Of the 18 stars, they were randomly assigned to one of 3 coolers (blocks); one cooler maintained constant temperature throughout the trial (11.5°C control), while the other two coolers held the elevated temperature (14.5°C) experiment. All stars had an additional 48h in these containers *prior* to treatment, which lasted 8 days. The unbalanced design was in expectation of increased mortality/symptomatic individuals in the elevated temperature experiment.

### Respirometry

Prior to temperature treatment but after acclimation to the constant base temperature of the OA lab, each individual was measured for VO_2_ as in Fly et al (2012), using a 10L chamber that excluded air pockets and contained a small 12V aquarium pump for mixing. The chamber itself was submerged in extra space in the lab coolers so that control or treatment temperatures were maintained. Into each chamber, an individual *P. ochraceus* was placed along with a pre-calibrated and initialized HOBO dissolved oxygen data logger (Onset model U26-001). The logger was allowed to equilibrate for 15 minutes and then data were collected for 45 minutes before the animal was returned to its individual chamber. VO_2_ was calculated as (Δ[O_2_] x *V*)/(*t* x *m*), *i.e*. the change in dissolved oxygen given the volume *V* of the chamber, the time *t* of analysis, and the mass *m* of the individual. Each individual was assayed prior to start of treatment temperatures and the last full day of the experiment.

## Results

### Sequencing of EF1A intron

We obtained 22 full-length sequences of the EF1A intron region (GenBank accession numbers MK092902-MK092924), including sequences representing both alleles at 9 of the 10 *P. ochraceus* individuals represented in Chandler and Wares (2017); the DNA from individual *Po*8 has degraded to the point that amplification of this region was not possible under these PCR conditions. Each individual had 2 recognizably distinct alleles; the sequence coverage for each allele ranged from 2-10x. With these data – which also include some of sequences obtained in Pankey & Wares (2009) we are able to re-assess the original hypothesis of homozygous lethality for the indel polymorphism. Sequence data for individuals from Pankey & Wares (2009) that clearly retained PCR-mediated recombination, e.g. failed the 4-gamete test among segregating sites, were excluded as that had been a concern in the original interpretation of the data (Pankey & Wares 2009). A neighbor-joining tree for these sequence data is shown in **Figure 1**; the rooting of the tree with congeners *P. brevispinus* and *P. giganteus* are particularly helpful in that the 6-bp “indel” region is found in both alleles from both congeneric species, indicating that the mutation observed in *P. ochraceus* is a deletion. These data indicate that of the 5 individuals previously inferred as “homozygous” for the deletion allele (allele ‘A’; designations in previous papers were based on ‘A’ being short and ‘B’ being long) using gel electrophoresis, there are two individuals (*Po*5, *Po*7) that each are homozygous for allele ‘B’ (with the 6-bp region), and three individuals (*Po*1, *Po*6) are homozygous for the deletion (capillary electrophoresis indicates that *Po*8 is homozygous for the deletion).

**Figure 1.**
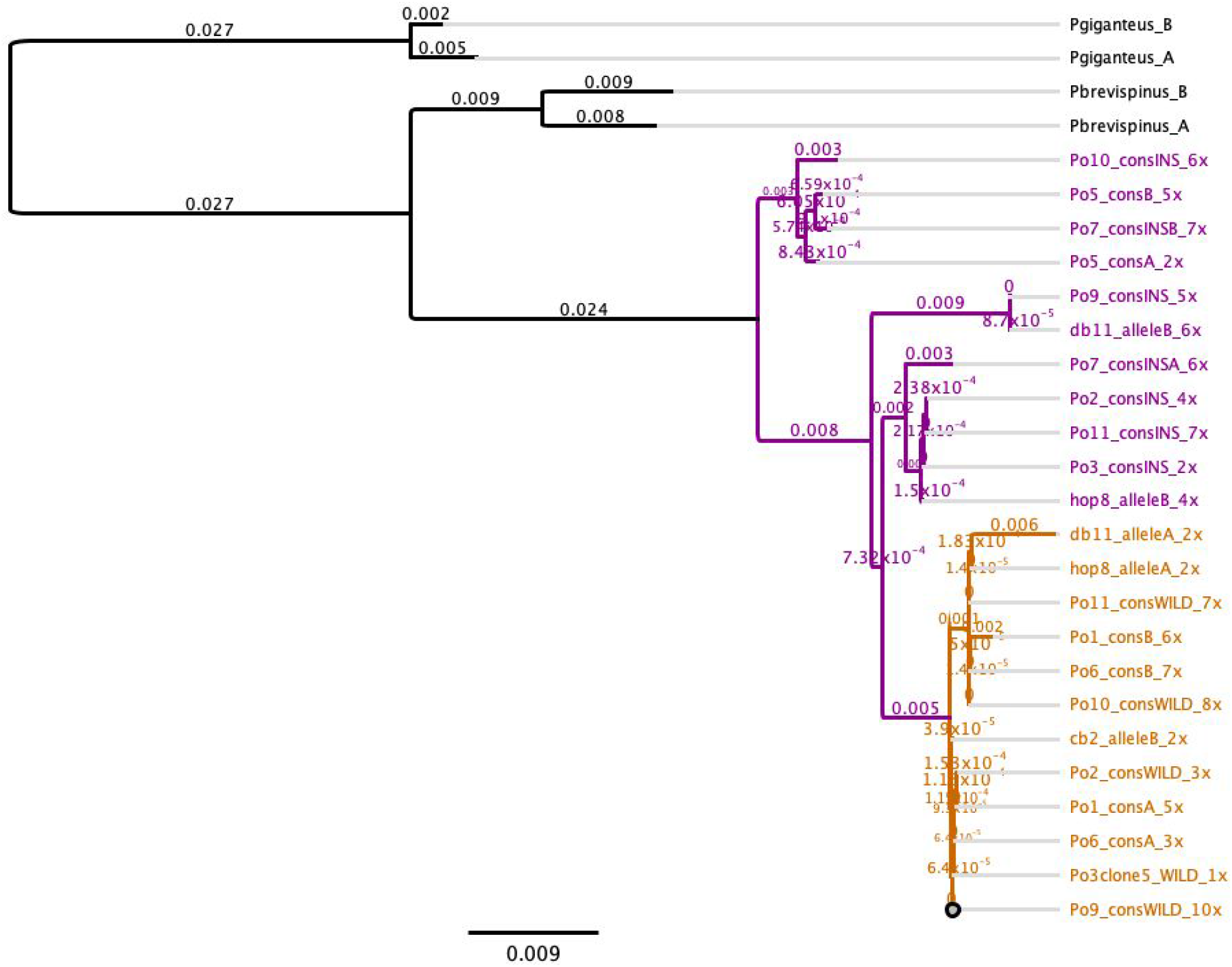
Congener-rooted gene tree for EF1A intron diversity in *P. ochraceus*. Alleles with the 6-bp focal region (allele class ‘B’) are shown in purple, alleles with deletion of this region are shown in orange (allele class ‘A’). Each allele is labeled with the individual identifier, information about which of 2 alleles, and the coverage for the consensus sequence of that allele.

From these data, we see much higher diversity in intron alleles containing the 6-bp region that separates the allele classes (*π* 0.00691) than in those alleles that do not have this 6-bp region (*π* 0.00076), with a higher number of segregating sites (4.857 versus 0.533), supporting the inference that the deletion event is derived and recent. Net nucleotide divergence of the two allele classes is 0.0097. Values of Tajima’s D test of neutrality are both positive (genus-typical *P. ochraceus* alleles 0.443, deletion alleles 1.303).

### Capillary electrophoresis of EF1A marker

The cloned sequences from *P. ochraceus* indicate that the 6-bp “insertion” from Pankey & Wares (2009) is not homozygous lethal. To clarify what is actually present, all the individuals from Chandler & Wares (2017) were re-amplified and a Fragment Analyzer automated capillary electrophoresis system (Advanced Analytical Technologies) was used to screen the amplicons. These results indicate that when the indel in question is heterozygous, a heteroduplex product is consistently generated that migrates more slowly (**Figure 2**). Thus, the appearance of two bands on an agarose gel as in Pankey & Wares (2009) or Wares & Schiebelhut (2016) is a reliable indicator that the individual is heterozygous, but the appearance of one band on an agarose gel is not an accurate indicator of homozygosity. Therefore, this same method was then used to re-genotype individual *P. ochraceus* from Wares & Schiebelhut (2016) and the individuals collected in 2018 for the respirometry work.

**Figure 2.**
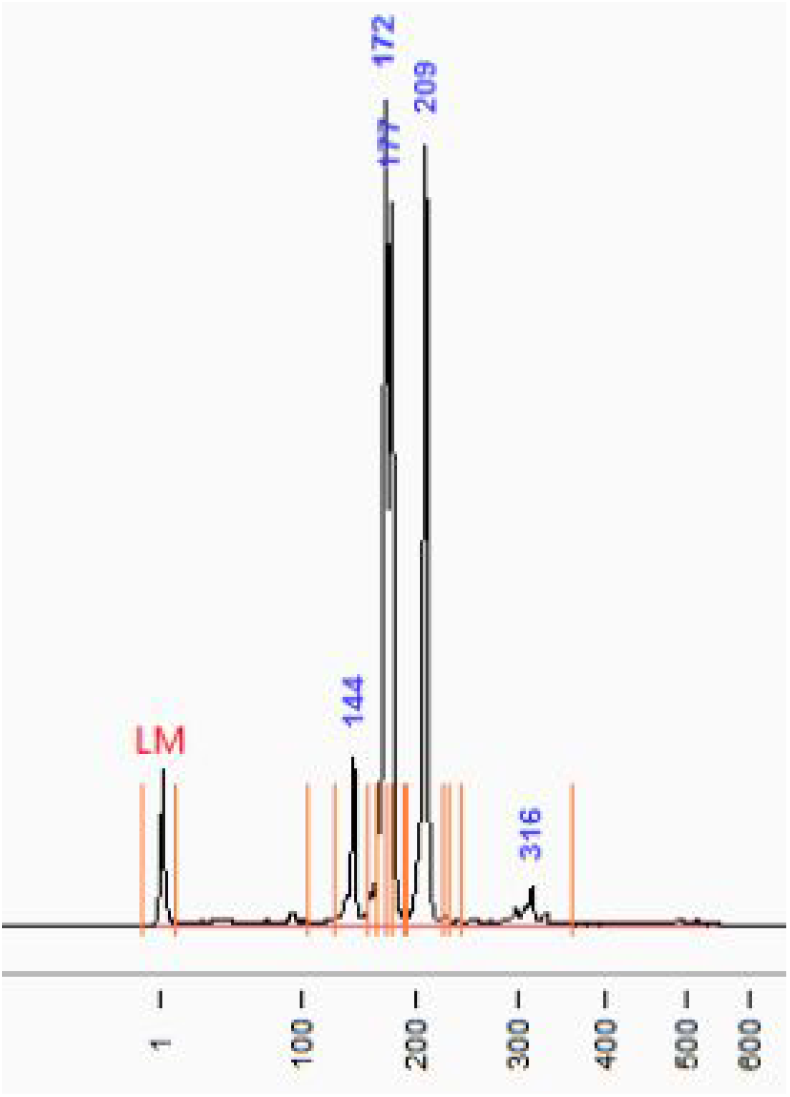
Example capillary electrophoresis of EF1A genotype reactions shows that individuals heterozygous for the 6-bp indel present both alleles at the correct sizes (172/177bp; the separation here is 5bp because of a linked 1bp deletion), as well as a heteroduplex product that migrates more slowly (indicated at 209bp).

Generally, we characterized individuals as carrying the “B” allele (lacking the 6-bp deletion) when there was a single peak present at ~173 bp (173.3 ± 1.7). The second major allele, “A,” was representative of the version of the intronic sequence with the 6-bp deletion. Peaks corresponding to allele “A” existed at ~179 bp (179.3 ± 1.3). Heterozygous individuals displayed peaks corresponding to alleles “A” and “B” that were consistently separated by ~6 bp (5.6 ± 0.7; a linked 1-bp deletion occasionally generated a 5-bp separation between peaks). Heterozygous genotypes were further distinguished by a PCR-generated heteroduplex product peak of similar magnitude at ~209 bp (209.5 ± 2.0; **Figure 2**).

Many of the individuals exhibited alleles with clear peak resolutions that were easily identifiable using the above criteria. Some, however, appeared to represent an intermediate between the two signatures. In these instances, where individuals consistently generated one prominent peak, but were accompanied by a second, smaller peak (both within the expected bp size range for a heterozygote), peak ratio rules were used to assign genotypes as either homozygous or heterozygous. More specifically, when one peak was greater than 5x the height of the other, the individual was designated as a homozygote. Only one of the eight individuals with questionable peak ratios fell below this mark, and was categorized as a heterozygote (“B” allelic peak was 2.9x the height of the “A” allelic peak).

Finally, one individual (FHLP39) generated consistent capillary electrophoretograms that were completely foreign to the set of objective rules described above. This individual was excluded from further analyses.

### Genotypic analysis

All individuals from the Wares & Schiebelhut (2016) collections were re-genotyped using capillary electrophoresis as basis for scoring individuals as homozygous for the 6-bp deletion (AA), heterozygous (AB), or homozygous without deletion (BB). Data are available in **Supplement 1**. Relative to the Wares & Schiebelhut data, 4 of 268 individuals were re-genotyped as one of the two homozygotes when previously scored as a heterozygote; 3 of 268 individuals were re-genotyped as a heterozygote when previously scored as a homozygote. Otherwise, of course the previous homozygotes were re-scored into one of two true homozygote classes, with resulting genotype frequencies of 0.24, 0.44, 0.32 respectively (there is a slight excess of homozygous individuals relative to Hardy-Weinberg expectations, but this is not statistically significant; p=0.183).

As in the prior analysis, the proportion of symptomatic heterozygotes (0.394; see **Figure 3**) is lower than the proportion of symptomatic homozygotes for this mutation in either genotype class (AA incidence of 0.527, BB incidence of 0.479). At any given location, at least one of the homozygous genotypes has a higher incidence of SSWD in these samples than the heterozygote genotypes. However, in contrast with the results of Wares & Schiebelhut (2016), binomial regression of these 3 *Pisaster* genotype classes against disease status indicates no relationship (p 0.412). When location of the collected individuals is included, the best model selected using AIC weighting included interactions between size, location, and genotype (AIC weight 0.906); in this GLM, the effect of genotype alone on disease status has improved relationship (p 0.10), with similar effects of location (p 0.11) and the interaction of genotype and location (p 0.10). Size itself had no predictive ability when tested alone (p 0.313) or in this interactive model (0.292).

**Figure 3.**
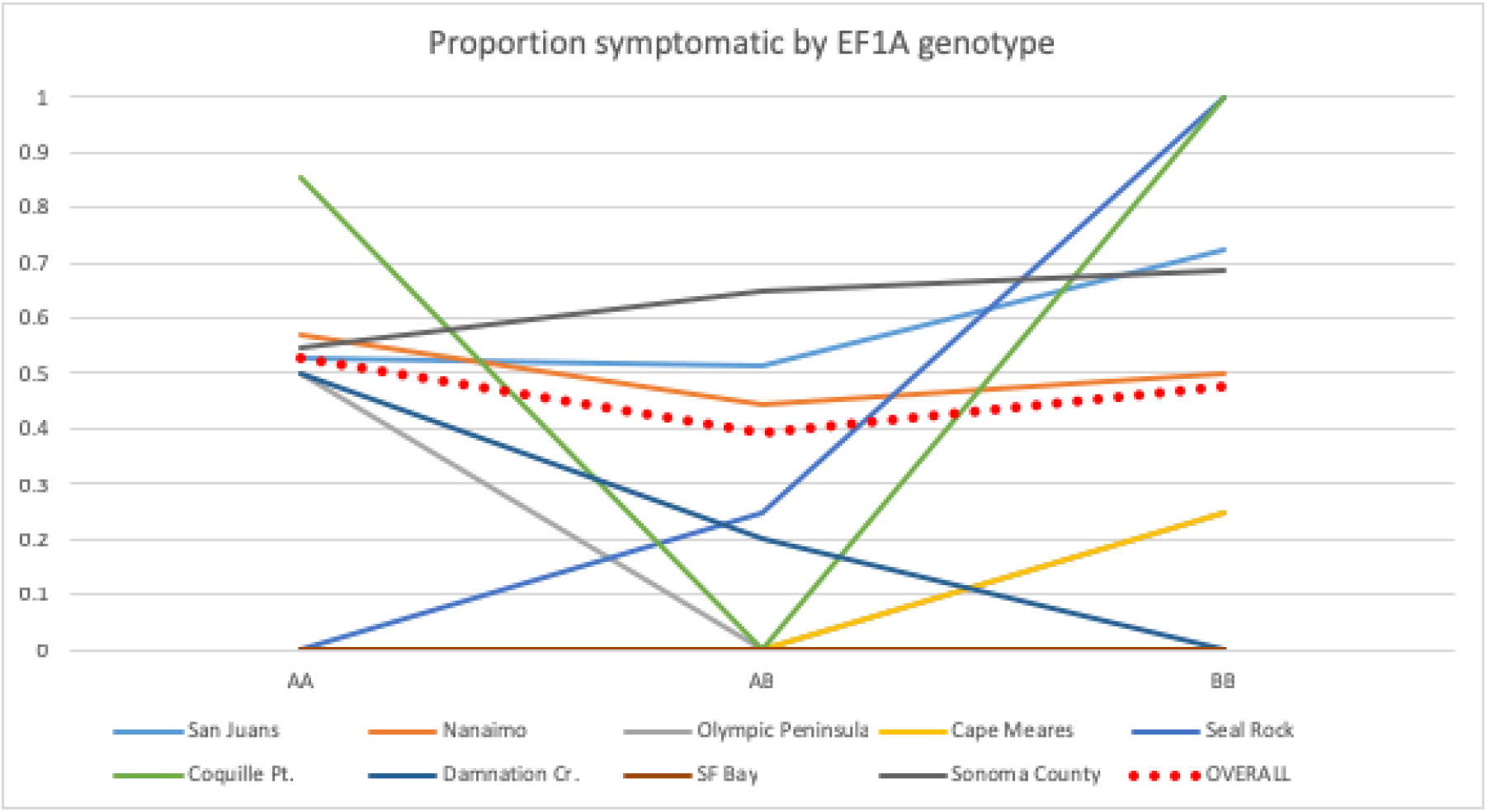
Proportion symptomatic plotted by pooled regional site (as in Wares & Schiebelhut 2016, e.g. collections within 10km of each other), and overall (red dotted plot). At all locations the incidence of symptomatic individuals is lower in heterozygous individuals than *at least one* class of homozygous individuals, but the difference is not statistically supported.

### Collection of individuals (2018) and Laboratory acclimation and experiment

The 18 individuals of *P. ochraceus* collected for the respirometry experiment are represented in **Table 1** with measured VO_2_ from before and after experimental treatment. The baseline “ambient” temperature for these trials was set at 11.5°C, the mean sea table water temperature measured over the previous 2 days under diurnal temperature fluctuation. Individuals were randomly assigned to treatment, with a larger number in the heat exposure trial in case of elevated mortality. Only a single individual showed symptoms of SSWD late in the trial; that individual subsequently began shedding arms and, after measurement of oxygen consumption, was frozen for further study.

**Table 1.** Individuals of *P. ochraceus* collected from field at same location on same date were weighed and fingerprinted for the EF1A marker (AA, one band; AB, two bands) prior to respirometry analysis. Individuals were subsequently genotyped using CE as noted; AA, homozygous deletion; AB, heterozygote; BB, homozygous no deletion). VO_2_ is presented following 2 days acclimation to lab conditions at constant 11.5°C, and following 7 days exposure to control (11.5°C) or elevated (14.5°C) temperatures. Individual Pis18_13, marked with an asterisk, exhibited signs of SSWD on the last day of exposure and the second respirometry measurement.

| Individual | Mass (g) | EF1A fingerprint (gel) | EF1A genotype (capillary) | VO<sub>2</sub> @ 11.5° | VO<sub>2</sub> after (treatment) |
| --- | --- | --- | --- | --- | --- |
| Pis18_01 | 235.75 | AA | AA | 0.023188 | 0.022057 (control) |
| Pis18_02 | 357.75 | AB | AB | 0.021989 | 0.022361 (heat) |
| Pis18_03 | 333.25 | AA | AA | 0.022005 | 0.020805 (control) |
| Pis18_04 | 110.75 | AB | AB | 0.036117 | 0.028893 (control) |
| Pis18_05 | 310.75 | AA | BB | 0.022311 | 0.025744 (heat) |
| Pis18_06 | 274.75 | AA | BB | 0.026691 | 0.022808 (control) |
| Pis18_07 | 106.25 | AB | AB | 0.031373 | 0.033882 (heat) |
| Pis18_08 | 123.25 | AA | AA | 0.021636 | 0.029208 (heat) |
| Pis18_09 | 139.25 | AB | AB | 0.037343 | 0.022023 (heat) |
| Pis18_10 | 89.25 | AA | BB | 0.016433 | 0.034360 (heat) |
| Pis18_11 | 95.75 | AB | AB | 0.030635 | 0.033420 (control) |
| Pis18_12 | 135.25 | AB | AB | 0.028589 | 0.028589 (control) |
| Pis18_13* | 403.75 | AB | AB | 0.020805 | 0.017172* (heat) |
| Pis18_14 | 257.75 | AA | BB | 0.019139 | 0.026899 (heat) |
| Pis18_15 | 159.25 | AB | AB | 0.025955 | 0.035164 (heat) |
| Pis18_16 | 218.25 | AA | AA | 0.022604 | 0.025658 (heat) |
| Pis18_17 | 187.25 | AB | AB | 0.022074 | 0.053404 (heat) |
| Pis18_18 | 381.75 | AB | AB | 0.020258 | 0.026195 (heat) |

### Respirometry

With individuals binned by agarose gel phenotype (indel heterozygous “AB” versus homozygous “AA”, **Table 1**), our analysis found evidence against the hypothesis that both genotype classes have equal measures of respiration (t-test, p=0.023) under ambient water temperature conditions (11.5°C). The mean VO_2_ for “homozygous” individuals (n=8) was 0.0217(+/− 0.0029), while the mean for heterozygous individuals (n=10) was 0.0275(+/− 0.0062). This effect size is large, with Cohen’s D of 0.997. Following elevated temperature treatment (control 11.5°C, treatment 14.5°C), our results are less easily interpreted: in the control treatment, the mean individual contrast in VO_2_ from the first measurement point to the second was negligible for heterozygotes (mean change −0.0015, n=3) and homozygotes (mean change −0.0021, n=3) with an overall test of control individuals indicating no distinction in the paired samples (paired t-test, p=0.261), while in the treatment chambers there was an expected increase in VO_2_ (heterozygotes mean change 0.0043, homozygotes mean change 0.0079) but the result was no distinction in VO_2_ among genotype classes (paired t-test, p=0.259). We also tested for differences in mass among the fingerprint classes and found no difference (p=0.652), and no effect of block/table (p=0.476).

When individuals are represented with their revised genotype (Table 1) the mean VO_2_ for individuals homozygous for the EF1A intron deletion (AA in Table 1) is 0.0224(+/− 0.0007), and the remaining “homozygous” individuals (BB in Table 1) is 0.0211 (+/− 0.0044). The sample size is now much smaller, but the comparison of VO_2_ between AB individuals and AA individuals indicates these distributions are not the same (p = 0.029) while the comparison between AB individuals and BB does not indicate they are drawn from different distributions (p=0.06; **Figure 4**). The effects of the temperature treatment itself are less easily assessed with very small size classes by genotype; the mean VO_2_ of AA individuals at the end of the trial was 0.027433(+/−0.0025, n=2) and for BB individuals was 0.0301(+/−0.0061, n=3), but as above there was no difference indicated between the genotype classes after heat exposure.

**Figure 4.**
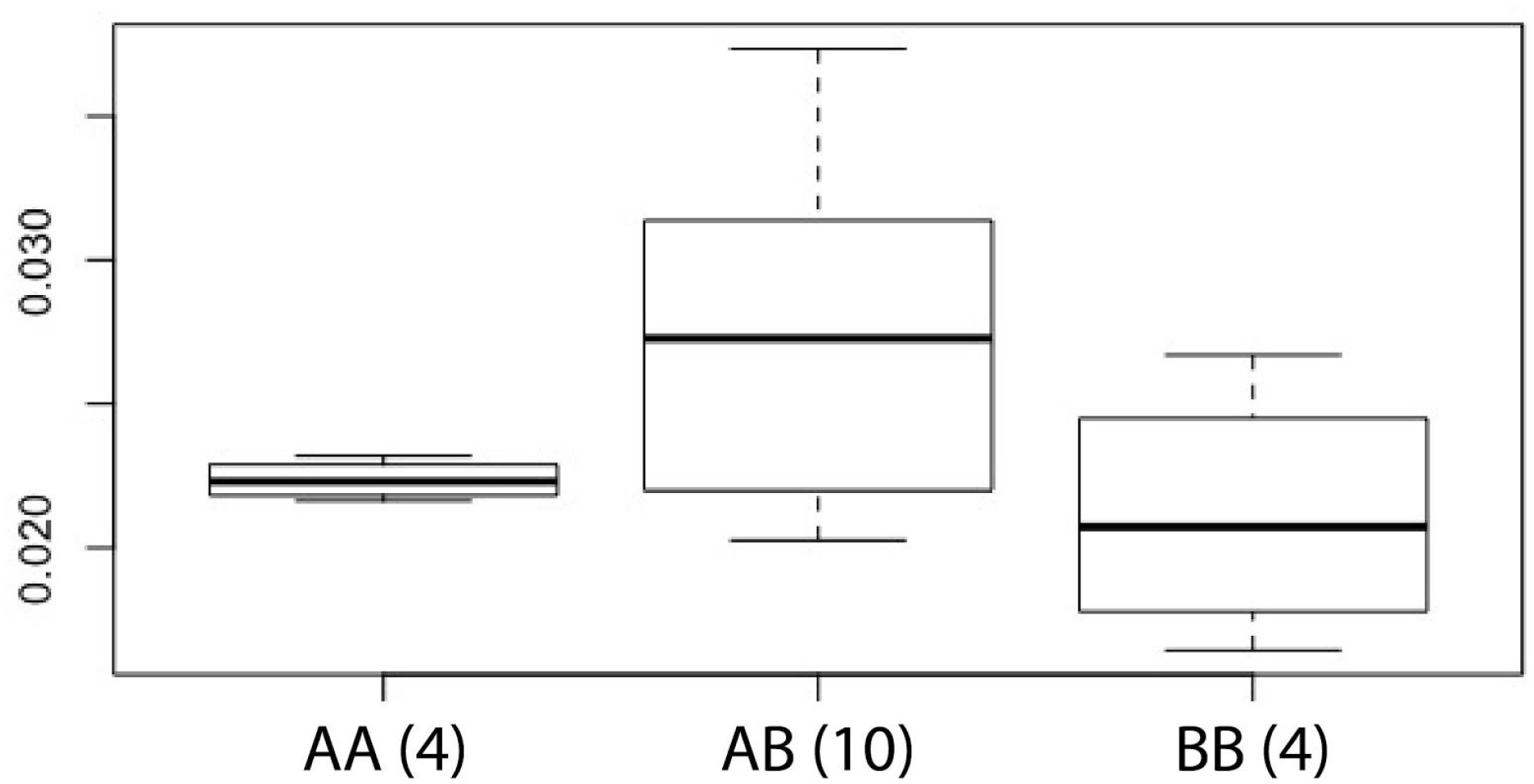
Comparison of VO_2_ measured for genotyped *P. ochraceus* (sample size of each shown on axis) at ambient (11.5°C) temperature following 2 days acclimation to lab conditions.

## Discussion

### Changing the genetic model

Sequence data obtained from high fidelity cloning under conditions meant to limit PCR-mediated recombination showed clearly that the indel mutation studied to date can be found in all 3 possible genotypes for the 2 allele classes. Though a small number of individuals were also cloned and sequenced both to develop and validate this marker (Pankey & Wares 2009), the EF1A intron fingerprint as seen on agarose was pursued as an apparent instance of confirmation bias: two electrophoretically distinct alleles were expected, and two electrophoretically distinct bands were observed. However, more thorough sequencing of cloned alleles and re-genotyping of individuals with capillary electrophoresis highlights this as a case of a heteroduplex (sometimes used to identify indel polymorphisms, in fact; Shore and Myerowitz 1990) being formed in the late cycles of PCR, rather than the genotypes being observed directly on the agarose gels (gel purification and Sanger sequencing of the “large” fragment only recovered the expected “small” sequences; JPW, pers. obs.). Reliance on this artifactual method to test the frequency of genotypes in offspring of crosses led to a misunderstanding about the nature of the focal mutation. Here we show that the 6-bp indel mutation can be found in heterozygous and both homozygous states, and the flanking sequence diversity is complex with no clear linked diversity that could otherwise have generated the ‘outlier’ signal of individual *Po*5 in Chandler & Wares (2017).

However, the ability to separate individuals of *P. ochraceus* into genotypic classes based on this mutation remains intriguing, in terms of fitness, transcriptional regulation, and physiology. Despite the falsified model underlying the analyses in Wares & Schiebelhut (2016), individuals heterozygous for this 6-bp mutation had an ecologically relevant increase in survival following the 2013-2014 outbreak of SSWD over individuals that were homozygous for either allele; this is still supported in our re-analysis of those data, but the revised genotypic model indicates no significant difference. Additionally, individuals heterozygous for the 6-bp mutation displayed notably distinct transcriptional profiles under ambient temperature conditions, and had only minor transcriptional responses to heat stress relative to individuals that were homozygous for either allele (Chandler & Wares 2017; **Figure 5**). Finally, in the current study, individuals that are heterozygous for the focal mutation have higher rates of oxygen consumption at ambient temperature conditions, and have a much lower change in respiratory rate with temperature stress, than individuals homozygous for either allele. This result in itself was predicted in part from differential righting responses in a subset of individuals from Chandler & Wares (2017). Thus, after re-evaluating the genotype inferences from the original study we continue to find biological signal to this DNA fingerprint that could be worth further pursuit.

**Figure 5.**
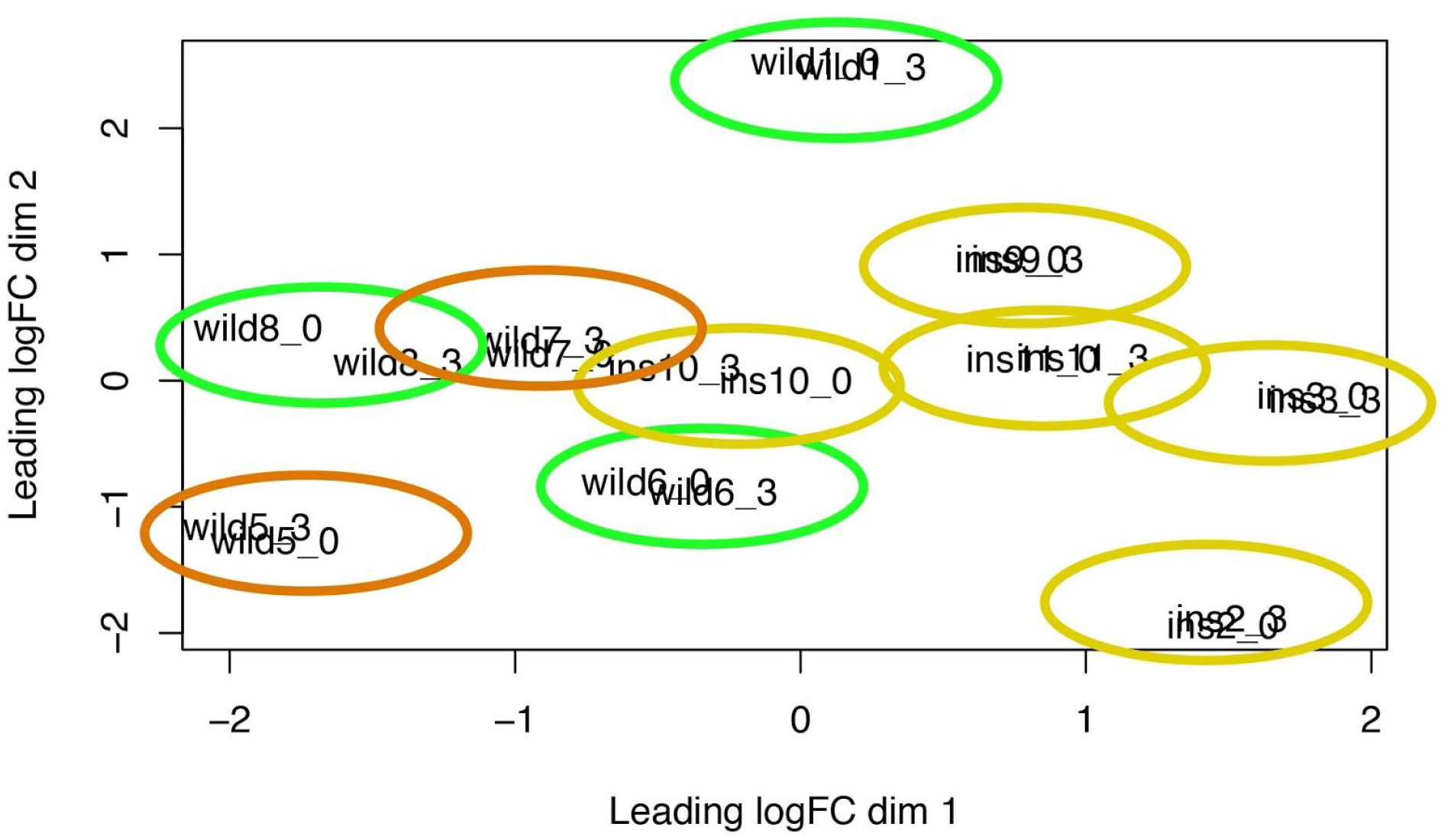
MDS plot reflecting differential expression of individuals by genotype, using the data from Chandler & Wares (2017). Each individual is labeled by electrophoretic EF1A phenotype class, individual ID, and treatment conditions (_0 indicating transcript profiles under ambient conditions, _3 indicating transcript profiles following heat exposure). EF1A genotypes as re-assessed are colored differently for contrast: light green individuals are AA, yellow are heterozygous AB, and orange individuals are BB. Reproduced from supplementary information in Chandler & Wares (2017)

### Is there differential fitness relative to SSWD among genotypes?

Re-genotyping the individuals from the Wares & Schiebelhut study continue to indicate lower incidence of SSWD in the heterozygous individuals for this mutation but with low effect size; there is not a clear signature of whether one homozygous genotype performs better than the other, though at sites with larger sample sizes the incidence of SSWD tends to be higher in the BB homozygotes. We note that another independent attempt to replicate these results (using the artifactual electrophoretic fingerprint) did not indicate differential incidence (S. Gravem, personal communication). However, in the initial study (Wares & Schiebelhut 2017) and in the current data, there is also a consistent signature of each location independently presenting higher incidence in at least one of the homozygote classes, and in all but one site higher incidence among all homozygotes, than in heterozygotes.

### Are there biological differences based on genotype?

As shown in **Figure 5**, RNA expression profiles are distinguishable among genotype classes under ambient conditions, suggesting the potential for constitutive differences in their biology. The differential gene expression results were robust to two forms of permutational tests (Chandler & Wares 2017), which is how individual *Po*5 (the individual that triggered this investigation) was highlighted as unusual in the first place. Unfortunately, we do not have identification of sex for these individuals; it is plausible that our genetic marker data actually reflect sex-based differences in biology (with no causal association), or other unrecognized traits. Re-genotyping these individuals clarifies our assessment that there is not a detectable difference in righting response times across all of the 2016 individuals (Chandler & Wares 2017), which had been used to evaluate activity levels that may differ among individuals both with and without temperature exposure.

In attempting a more refined approach to this latter question – in an experiment designed before recognition of the genotype/fingerprint artifact – we replicated the 2016 experiment in a more controlled lab environment and evaluated the respiratory physiology of individuals at ambient and elevated temperature exposure. Though our sample size is limited, there appears to be a signature for EF1A-heterozygous individuals having higher baseline oxygen consumption (**Figure 4**) and a more modest response to temperature exposure, similar to the reduced transcriptional response that fingerprint-based heterozygotes exhibited in the previous temperature exposure work (Chandler & Wares 2017).

### Conclusions

An intriguing result that supported the idea that genomic diversity may benefit populations faced with disease challenges (e.g., Brock et al. 2015) appears now to be less clearly interpretable. We note that the likely origins of this misunderstanding about the inheritance mode of mutational diversity in this EF1A intron can be categorized as a form of confirmation bias (Nickerson 1998) paired with a failure to recognize that not rejecting an alternative hypothesis is not the same as determining that hypothesis to be true (Gerrodette 2011). The initial work reflected in Pankey & Wares (2009) was not specifically funded research, and as current funding for understanding disease phenomena in *P. ochraceus* was not available at the time, two recent papers (Wares & Schiebelhut 2016, Chandler & Wares 2017) relied on the earlier work as a basis for exploration. One of our first priorities with funding, then, was to assess our underlying assumptions thoroughly before progressing further with the ‘homozygous lethal’ model for the focal mutation in this paper; we have succeeded in showing that model to be incorrect.

Though there are still biological signals reflecting the likelihood of ecologically relevant phenotypic diversity in *P. ochraceus*, the original intent of exploring the marker was as a simple and inexpensive evaluation of diversity at a nuclear marker that would also satisfy reviewers. Subsequent explorations were pursued both because of the original interpretation (Pankey & Wares 2009) and the ease and cost of testing this marker against SSWD prevalence. Given the sequence complexity of this EF1A intron region and the additional technical effort required to characterize this diversity for population-level analysis, we recognize that our update suggests the costs and benefits of further study are now very different.

## Acknowledgments

Initial feedback, and in particular kind reassurance, was greatly appreciated from Allen Moore, Erik Sotka, Morgan Kelly, Sarah Gravem, Morgan Eisenlord, Mike Hart, Mike Dawson, Lauren Schiebelhut, Sara Heisel, and others. Thanks go to Amy Rosemond, Philip Bumpers, Catherine Sweere, and Ben Miner for helping us develop the application of “pecan bags” for experimental tagging – a notable advance in manipulative work with *Pisaster*. Mike Dawson, Dannise Ramos, Lauren Schiebelhut, and Ian Hewson contributed greatly to initial and modified experimental design. Many thanks to Noah Workman and others at the Georgia Genomics and Bioinformatics Core, with genuine respect for the contributions of the late Jeff Wagner. The staff and scientists at the Friday Harbor Laboratory, in particular Becca Guenther, make this work possible and enjoyable. This manuscript was greatly improved following comments from Mike Dawson and Sabrina Pankey. Funding for this work comes from NSF-OCE award 1737381 to JPW.

